# Combining mutation and recombination statistics to infer clonal families in antibody repertoires

**DOI:** 10.1101/2022.12.22.521661

**Authors:** Natanael Spisak, Thomas Dupic, Thierry Mora, Aleksandra M. Walczak

## Abstract

B-cell repertoires are characterized by a diverse set of receptors of distinct specificities generated through two processes of somatic diversification: V(D)J recombination and somatic hypermutations. B cell clonal families stem from the same V(D)J recombination event, but differ in their hypermutations. Clonal families identification is key to understanding B-cell repertoire function, evolution and dynamics. We present HILARy (High-precision Inference of Lineages in Antibody Repertoires), an efficient, fast and precise method to identify clonal families from high-throughput sequencing datasets. HILARy combines probabilistic models that capture the receptor generation and selection statistics with adapted clustering methods to achieve consistently high inference accuracy. It automatically leverages the phylogenetic signal of shared mutations in difficult repertoire subsets. Exploiting the high sensitivity of the method, we find the statistics of evolutionary properties such as the site frequency spectrum and *d_N_/d_S_* ratio do not depend on the junction length. We also identify a broad range of selection pressures scanning two orders of magnitude.

## I. INTRODUCTION

B cells play a key role in the adaptive immune response through their diverse repertoire of immunoglobulins (Ig). These proteins recognize foreign pathogens in their membrane-bound form (called B-cell receptor or BCR), and battle them in their soluble form (antibody). Each B cell expresses a unique BCR that can bind their antigenic targets with high affinity. The set of distinct BCR harbored by the organism is highly diverse [1], thanks to two processes of diversification: V(D)J recombination and somatic hypermutation. These stochastic processes ensure that repertoires can match a variety of potential threats, including proteins of bacterial and viral origin that have never been encountered before.

V(D)J recombination takes place during B cell differentiation [2, 3]. For each Ig chain, V, D, and J gene segments for the heavy chain, and V and J gene segments for the light chain, are randomly chosen and joined with random non-templated deletions and insertions at the junction, creating a long, hypervariable region, called the Complementarity Determining Region 3 (CDR3) (Fig. 1A). Cells are subsequently selected for the binding properties of their receptors and against autoreactivity. At this stage, the repertoire already covers a wide range of specificities. In response to antigenic stimuli, B cells with the relevant specificities are recruited to germinal centers, where they proliferate and their Ig-coding genes undergo somatic hypermutation [4] in the process of affinity maturation. Somatic hypermutation consists primarily of point substitutions, as well as insertions and deletions, restricted to Ig-coding genes [5]. The mutants are selected for high affinity to the particular antigenic target, and the best binders further differentiate into plasma cells and produce high-affinity antibodies. A more diverse pool of variants forms the memory repertoire, leaving an imprint of the immune response that can be recalled upon repeated stimulation.

**FIG. 1.**
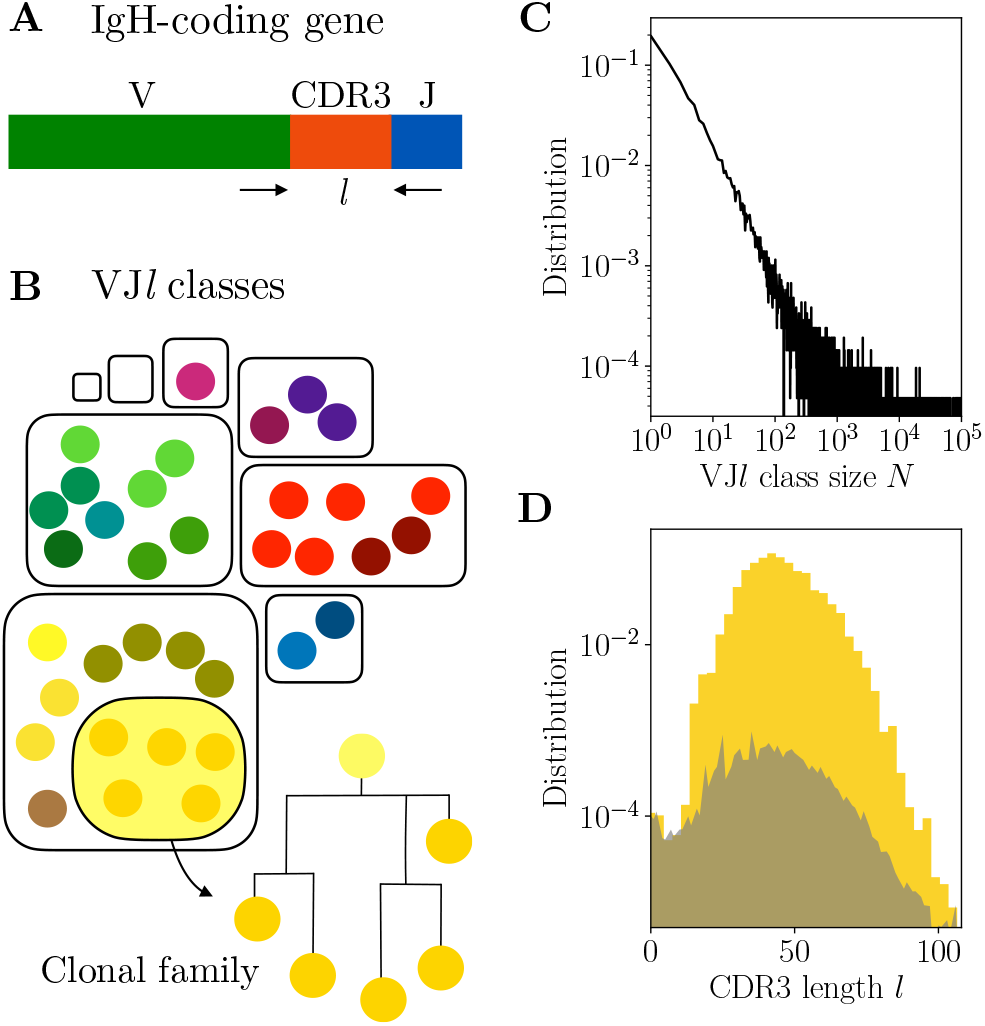
Clonal families and VJ*l* classes. (A) Variable region of the IgH-coding gene. (B) A clonal family is a lineage of related B cells stemming from the same VDJ recombination event. The partition of the BCR repertoire into clonal families is a refinement of the partition into VJ*l* classes, defined by sequences with the same V and J usage and the same CDR3 length *l*. (C-D) Properties of VJ*l* classes in donor 326651 from [1]. (C) Distribution of VJ*l* class sizes exhibits powerlaw scaling. The total number of pairwise comparisons in the largest VJ*l* classes is ~ 10^52^ = 10^10^. (D) Distribution of the CDR3 length *l*. The distribution is in yellow for in-frame CDR3 sequences (*l* multiple of 3), and in gray for out-of-frame sequences.

A clonal family is defined as a collection of cells that stem from a unique V(D)J rearrangement, and has diversified as a result of hypermutation, forming a lineage (Fig. 1B). These families are the main building blocks of the repertoire. Since members of the same family usually share their specificities [6], affinity maturation first competes families against each other for antigen binding in the early stages of the reaction, and then selects out the best binders within families in the later stages [7, 8].

High-throughput sequencing of single receptor chains offers unprecedented insight into the diversity and dynamics of the repertoire. Recent experiments have sampled the repertoires of the immunoglobulin heavy chain (IgH) of healthy individuals at great depth to reveal their structure [1]. Disease-specific cohorts are now routinely subject to repertoire sequencing studies, which help to quantify and understand the dynamics of the B-cell response [9, 10].

Partitioning BCR repertoire sequence datasets into clonal families is a critical step in understanding the architecture of each sample and interpreting the results. Identifying these lineages allows for quantifying selection [11–13] and for detecting changes in longitudinal measurements [10, 14]. In recent years, many strategies have been developed that take advantage of CDR3 hypervariability [15]: it is generally unlikely that the same or a similar CDR3 sequence be generated independently multiple times [13, 16]. Other approaches make use of the information encoded in the intra-lineage patterns of divergence due to mutations [17, 18]. All inference techniques need to balance accuracy and speed. Simpler methods are fast but have low precision, while more complex algorithms have long computation times that do not scale well with the number of sequences. This prohibits the analysis of recent large-scale data such as [1].

In this work, we propose a new method for inferring clonal families from high-throughput sequencing data that is both fast and accurate. We use probabilistic models of junctional diversity to estimate the level of clonality in repertoire subsets, allowing us to tune the sensitivity threshold *a priori* to achieve a desired accuracy. We have developed two complementary algorithms. The first uses a very fast CDR3-based approach that avoids pairwise comparisons, while the second additionally exploits information encoded in the phylogenetic signal outside of the junction. We compare our method with state-of-the-art approaches in a benchmark with realistic synthetic data.

## II. RESULTS

### A. Analysis of pairwise distances within VJ*l* classes

A common strategy for partitioning a BCR repertoire dataset into clonal families is to go through all pairs of sequences and identify pairs of clonally related sequences. In the following, we call such related pairs *positive*, and pairs of sequences belonging to different families *negative*. Then, the partition is built by single-linkage clustering, which consists in recursively grouping all positive pairs. Two characteristics of the repertoire complicate the search for this partition: large total number of pairs and low proportion of positive pairs, called *prevalence* and denoted by *ρ*. Low prevalence is usually due to a high frequency of singletons, which are sequences with no relative in the dataset.

A pair of related sequences is expected to share the same V and J gene usage, as well as the same CDR3 length *l*, determined by alignment to the templates (Figure 1A). For a description of the data preprocessing and alignment to the V and J gene templates see Methods section IV A.

The methods developed here begin by partitioning the data into VJ*l* classes, defined as subsets of sequences with the same V and J gene usage, and CDR3 length *l* (Fig. 1B). Clonal relationships are then found within these classes. While this severely limits the number of unnecessary comparisons, some VJ*l* classes still exceed 10^5^ sequences in large datasets, leading to the order of 10^10^ pairs (see Figure 1C for the distribution of the VJ*l* class sizes *N* for donor 326651 of [1]).

The signature of the VDJ rearrangement is largely encoded by the CDR3 alone. As we will see, CDR3 length has a strong impact on the difficulty of clonal family reconstruction. The distribution of CDR3 lengths *l* observed in the data is shown in Fig. 1D. In what follows we focus primarily on productive sequences, defined by their CDR3 length being a multiple of 3, which dominate most sequencing datasets. In addition, we restrict our analysis to CDR3 lengths between 15 and 105: shorter or longer junctions have comparable frequencies to nonproductive junctions, suggesting that they are nonfunctional.

Within each VJ*l* class, we compute the Hamming distance *n* of each pair of CDR3s, defined as the number of positions at which the two nucleotide sequences differ. The distribution of these distances normalized by the CDR3 length, denoted by *x*, shows a clear bimodal structure, with two identifiable components (Fig. 2A): the contribution of positive pairs (of proportion *ρ*) peaks near *x* = 0 and decays quickly, whereas the bell-shaped contribution of negative pairs (of proportion 1 – *ρ*) peaks around *x* = 1/2.

**FIG. 2.**
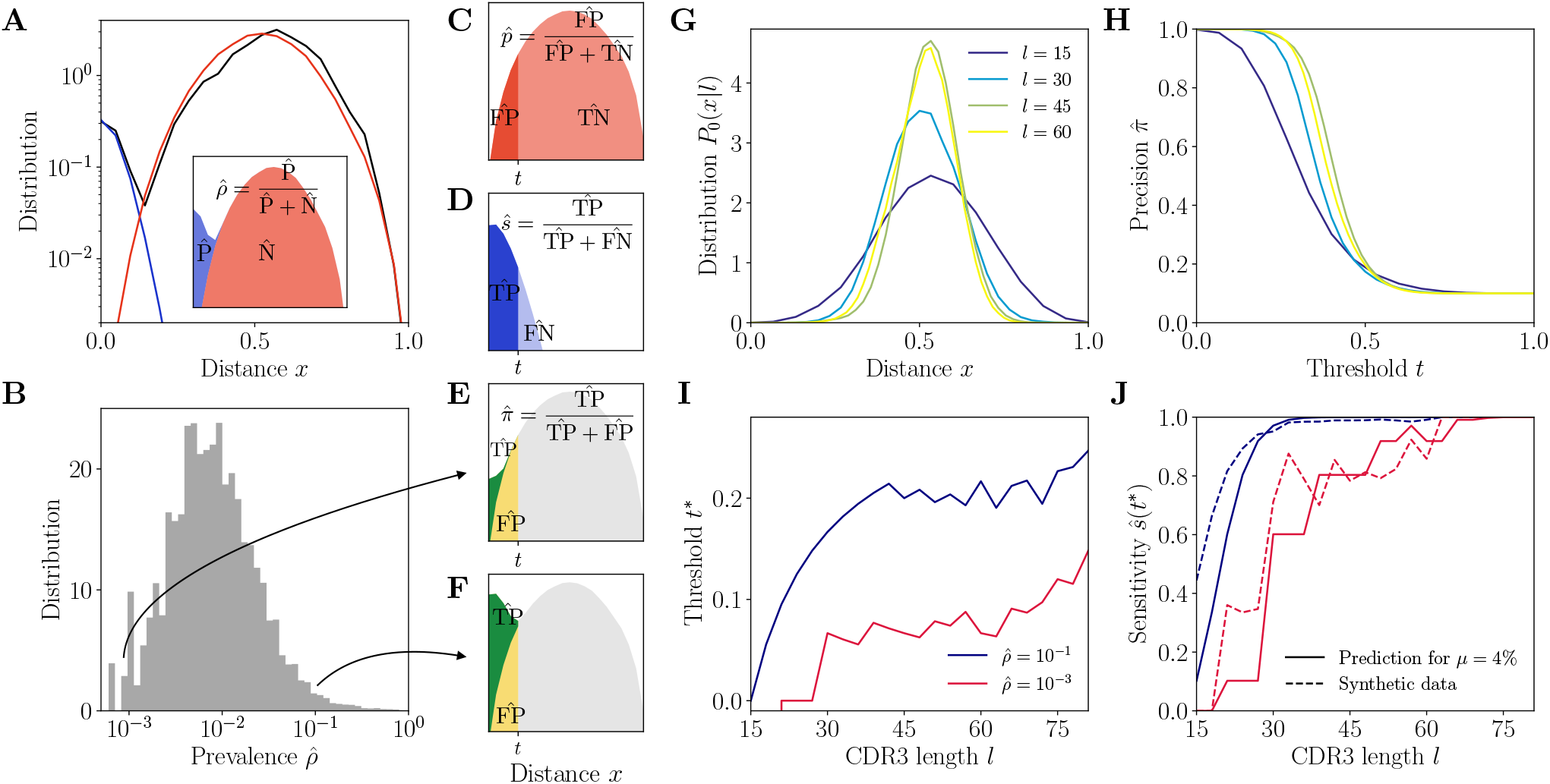
CDR3-based inference method. (A) Example distribution of normalized Hamming distances, *x* = *n/l*, for one VJ*l* class with CDR3 length *l* = 21, V gene IGHV3-9 and J gene IGHJ4 (black). We fit the distribution by a mixture of positive pairs (belonging to the same family, in blue), and negative pairs (belonging to different families, in red). See Figure S5 for example fit results across different CDR3 lengths. Inset: the prevalence is defined as a fraction of positive pairs and was estimated to 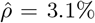. Data from donor 326651 of [1]. (B) Distribution of the maximum likelihood estimates of prevalence 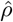 across VJ*l* classes. (C-F) The choice of threshold *t* on the normalized Hamming distance *x* translates to the following *a priori* characteristics of inference. (C) Fallout rate 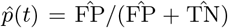. The null distribution of all negatives (N=FP+TN) is estimated using the soNNia sequence generation software. (D) Sensitivity 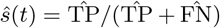. (E-F) Precision 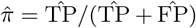. For the same choice of threshold *t*, a low prevalence of 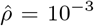 (E) leads to lower precision than high prevalence of 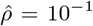 (F). (G) Null distribution *P*_0_(*x|l*) of distances between unrelated sequences, for *l* = 15, 30, 45, 60, computed by the soNNia software. (H) Precision 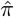 as a function of the threshold *t*, for *l* = 15, 30,45, 60 (colors as in G). (I) High-precision threshold *t*\* ensuring 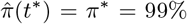, as a function of CDR3 length *l* for different values of the prevalence 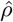. (J) Sensitivity *ŝ*(*t*\*) at the high-precision threshold *t*\*, as a function of CDR3 length *l* for different values of the prevalence 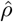 (colors as in I). Solid lines denote prediction for intermediate mean distance *μ* = 4%, dashed lines denote measurement in a synthetic dataset.

The prevalence *ρ* can be formally written as [∑_*i*_ *z_i_*(*z_i_* – 1)/2]/[*N*(*N* – 1)/2], where *z_i_* denote the sizes of the clonal families in the VJ*l* class, but we do not know these sizes before the partition into families is found. To overcome this issue, we developed a method to estimate *ρ a priori*, without knowing the family structure (Methods section IV C). We do this by fitting the distribution of *x* as 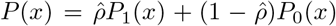, where *P*_1_(*x*) and *P*_0_(*x*) are the distributions of distances between positive and negative pairs (Fig. 2C and D), estimated as follows. *P*_0_(*x*) = *P*_0_(*x|l*) is computed for each length *l* by generating a large number of unrelated, same-length sequences with the soNNia model of recombination and selection [19], and calculating the distribution of their pairwise distances (Methods section IV B). *P*_1_(*x*) is approximated by a Poisson distribution, *P*_1_(*x*) = (*μl*)^*xl*^*e*^−*μl*^/(*xl*)!, with adjustable parameter *μ*. Fitting is performed with an expectation-maximization algorithm which finds maximum-likelihood estimates of the prevalence 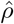 and mean intra-family distance 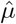 for each VJ*l* class.

The results of the fit show that 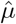 varies little between VJ*l* classes, around 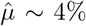 (Fig. S1). In contrast, the prevalence 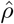 varies widely across classes (Fig. 2B), spanning three orders of magnitude. In addition, when we examine the VJ*l* classes with increasing CDR3 length *l*, we find that the positive mode *P*_1_(*x*) of the distribution varies little, whereas the negative mode *P*_0_(*x*) becomes more and more peaked around 1/2 (Fig. 2G and Fig. S2), making the two modes more easily separable.

### B. CDR3-based inference method with adaptive threshold

We want to build a classifier between positive and negative pairs using the normalized distance *x* alone, by setting a threshold *t* so that pairs are called positive if *x* ≤ *t*, and negative otherwise. Using our model for *P*(*x*), for any given *t* we can evaluate the number of true positives 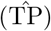 and false negatives 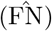 among all positive pairs 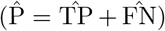, as well as true negatives 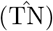 and false positives 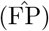 among the negative pairs 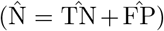, as schematized in Figs. 2C and D.

Our goal is to set a threshold *t* that ensures a high precision 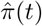, defined as a proportion of true positives among all pairs classified as positive (Fig. 2E). In a single-linkage clustering approach, we will join two clusters with at least one pair of positive sequences between them. Therefore, it is critical to limit the number of false positives, which can cause the erroneous merger of large clusters. We can write:

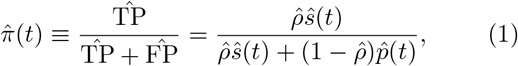

where 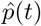 is the estimate of the fall-out rate (Fig. 2C), evaluated the from soNNia-computed *P*_0_:

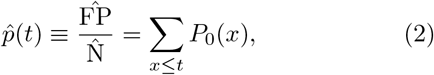

and *ŝ*(*t*) is the estimated sensitivity (Fig. 2D), evaluated from the Poisson fit to *P*_1_ (Methods, section IV D):

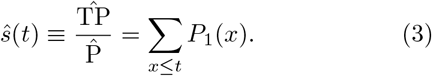

Finally, the estimated prevalence 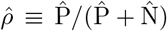 is inferred from the *P*(*x*) distribution as explained above.

Fig. 2H shows 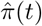 as a function of *t* for different CDR3 lengths and a fixed value of 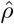. For each VJ*l* class, we define the threshold *t* = *t*\* that reaches 99% precision, 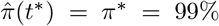, by inverting (1). This adaptive threshold depends on the VJ*l* class through the CDR3 length *l* and the prevalence *ρ*, and it increases with both (Fig. 2I): low clonality (small *ρ*) means few positive pairs and a smaller adaptive threshold, while short CDR3 means less information and a stricter inclusion criterion.

The resulting sensitivity, *ŝ*(*t*\*), which tells us how much of the positives we are capturing, is shown in Fig. 2J. We conclude that for a wide range of parameters, the method is predicted to achieve both high precision and high sensitivity. However, it is expected to fail when the prevalence and the CDR3 length are both low. At the extreme, for small values *ρ* and *l*, even joining together identical CDR3s (*t* = 0) results in poor precision because of convergent recombination (reflected by *t*\* < 0).

### C. Tests on synthetic datasets

So far we have presented a method to set a high-precision threshold with predictable sensitivity, based on estimates from the distribution of distances *P*(*x*) only. To verify that these performance predictions hold in real inference tasks, we designed a method to generate realistic synthetic datasets where the clonal family structure is known. This generative method will also be used in the next sections to create a benchmark for comparing different clustering algorithms.

Briefly, we first estimated the distribution of clonal family sizes from the data of [1] by applying the CDR3-based clustering method with adaptive threshold described above to VJ*l* classes for which the inference was highly reliable, i.e. for which the predicted sensitivity was *ŝ*(*t*\*) ≥ 90%. In that limit, clusters are clearly separated and the partition should depend only weakly on the choice of clustering method. The resulting distribution of clone sizes follows a power-law with exponent −2.3. Then, for each lineage, we draw a random progenitor using the soNNia model for IgH generation (Fig. S4), as well as a random size from the power-law distribution. Mutations are then randomly drawn on each sequence of the lineage in a way that preserves the mutation sharing patterns observed in families of comparable size from the partitioned data (Fig. S5). We thus generated 10^4^ lineages and 2.5 · 10^4^ sequences. More details about the procedure are given in the Methods, section IV E.

We applied the CDR3-based clustering method to this synthetic dataset. The sensitivity achieved at *t*\* roughly follows and sometimes even outperforms the predicted one *ŝ*(*t*\*) across different values of *ρ* and *l* (Fig. 2J, dashed line), validating the approach and the choice of the adaptive high-precision threshold *t*\* (the discrepancy is due to the fact that *μ* is assumed to be constant in the prediction, while it varies in the dataset). These results also confirm the poor performance of the method at low prevalences and short CDR3s.

### D. Incorporating phylogenetic signal

To improve the performance of the CDR3-based method, we set out to include the phylogenetic signal encoded in the mutation spectrum of the templated regions of the sequences. Two sequences belonging to the same lineage are expected to share some part of the mutational histories, and therefore sequences with shared mutations are more likely to be in the same lineage.

We focus on the template-aligned region of the sequence outside of the CDR3, where we can reliably identify substitutions with respect to the germline. We denote the length of this alignment by *L*, so that the total length of the sequence is *l*+*L*. For each pair of sequences, we define *n*_1_, *n*_2_ as the number of mutations along the templated alignment in the two sequences, *n*_0_ the number of mutations shared by the two, and *n_L_* = *n*_1_ + *n*_2_ – 2*n*_0_ the number of non-shared mutations. Under the hypothesis of shared ancestry, the *n*_0_ shared mutations fall on the shared part of the phylogeny, and are expected to be more numerous than under the null hypothesis of independent sequences, where they are a result of random co-occurrence (Fig. 3A).

**FIG. 3.**
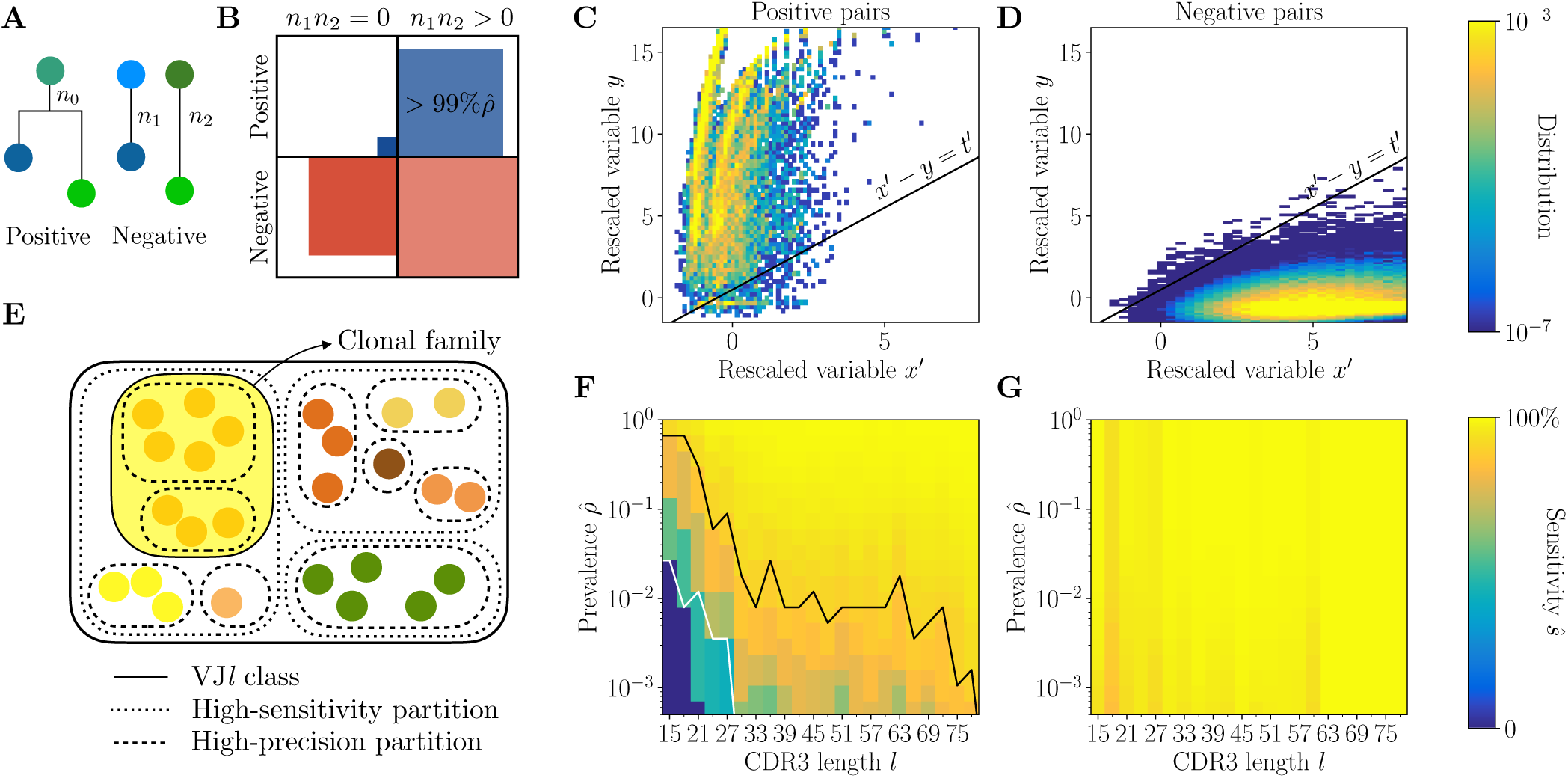
Full inference method with mutational information. (A) For a pair of sequences, *n*_1_, *n*_2_ denote the numbers of mutations along the templated region (V and J), and *n*_0_ is the number of shared mutations. For related sequences, *n*_0_ corresponds to mutations on the initial branch of the tree, and is expected to be larger than for unrelated sequences, where *n*_0_ corresponds to coincidental mutations. (B) Positive and negative pairs are called mutated if both sequences have mutations *n*_1_, *n*_2_ > 0. Among positive pairs in the synthetic datasets, more than 99% are mutated. (C, D) Distributions of the rescaled variables *x′* and *y* (4), for pairs of synthetic sequences belonging to the same lineage (positive pairs) and sequences belonging to different lineages (negative pairs). The separatrix *x′* – *y* = *t′* marks a high-precision (99%) threshold choice. (E) To limit the number of pairwise comparisons we make use of high-precision and high-sensitivity CDR3-based partitions. High precision corresponds to the choice *t* = *t*\*. High sensitivity corresponds to a coarser partition where *t* is set to achieve 90% sensitivity. When the two partitions disagree, mutational information can be used to break the coarse, high-sensitivity partition into smaller clonal families. (F,G) Mutations-based methods achieve high sensitivity across all CDR3 lengths *l* in the synthetic dataset (G), extending the range of applicability with respect to the CDR3-based method (F).

To balance the tradeoff between the information encoded in the templated part of the sequence and the recombination junction, we can compute characteristic scales for the two variables of interest: the number of shared mutations *n*_0_ and the CDR3 divergence *n*. Intuitively, in highly mutated sequences, we can expect substantial divergence in the CDR3. At the same time, the number of mutations in the templated regions would increase, possibly leading to more shared mutations. Conversely, sequences with few or no mutations carry no information in the templated region, but we also expect their CDR3 sequences to be nearly identical. To adapt a clustering threshold to the two variables, we compute their expectations under the two assumptions, and define the rescaled variables

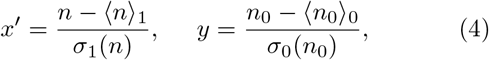

where 〈*n*_1_〉_1_ = *l*(*n_L_* + 1)/*L* is the expected value of *n*_1_ under the hypothesis that sequences belong to the same lineage, and 〈*n*_0_〉_0_ = *n*_1_*n*_2_/*L* is the expected value of *n*_0_ under the hypothesis that they do not. The standard deviations are likewise defined as 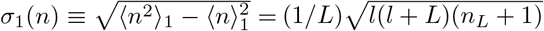 and 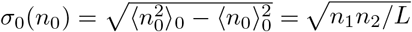 (Methods, section IVF).

For more than 99%of positive pairs, both sequences are mutated, i.e. *n*_1_, *n*_2_ > 0 (Fig. 3B). Without loss of sensitivity, we focus on the mutated part of the dataset, since we cannot use *y* for non-mutated sequences. The distributions of *x′* and *y* for positive and negative pairs (Figs. 3C and D) are well separated, with positive pairs characterized by an overrepresentation of shared mutations. By adding the phylogenetic signal *y* we can identify positive pairs of sequences that have significantly diverged in their CDR3 (*x′* > 0) but share significantly more mutations than expected (large *y*).

Computing *y* for each pair of sequences is computationally expansive. To avoid examining all pairs, we first perform a “coarse” clustering of each VJ*l* class using the CDR3-based method with a high threshold *t* that ensures high sensitivity *ŝ* = 90%. Most of related sequences are grouped together, resulting in a high-sensitivity partition (Methods, section IV D and Fig. 3E). For sufficiently large *ρ* and *l*, this partition coincides with the high-precision partition defined above by 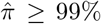. In that case, the CDR3-based method achieves good sensitivity and precision, and we do not need to compute *y*. When they do not coincide, we can use the phylogenetic signal *y* to refine the high-sensitivity partition. We only need to compute *y* for pairs that belong to the same coarse clusters: the phylogenetic signal is used to further break up these clusters into clonal families. This allows us to considerably reduce the number of pairwise comparisons that we need to make between the templated regions of the sequences.

Using *x′* and *y*, we classify pairs of sequences as positive (i.e. belonging to the same family) if *y* ≥ *x′* – *t′*, and as negative otherwise, where *t′* is a threshold chosen to achieve 99% precision in each VJ*l* class (Methods section IV F).

We then compute the expected sensitivity on the synthetic data, and find that it reaches values ≥ 90% across the whole range of prevalence *ρ* and CDR3 lengths *l*, outperforming the CDR3-based method in the low-*ρ*, low-*l* region (Fig. 3F and G). This proves that using the phylogenetic signal significantly improves performance over the CDR3-based method.

### E. Benchmark of the methods

We compare our approach to state-of-the-art methods, using the synthetic data described above as ground truth. In addition to our two algorithms—the CDR3-based method and the full method using both CDR3 and shared mutations—our benchmark includes the alignment-free method of [20], Partis [21], and Scoper [22].

First, we measure the inference time of each algorithm. We find that the inference time is primarily affected by the size of the largest VJ*l* class. Therefore, we measure the inference time using the largest class found in donor 326651 of dataset [1] with the size of *N* = 1.2 ×10^5^ unique sequences. We then apply the methods to a series of subsamples of this class to get the computational time as a function of the subsample size (Fig. 4A). We only allowed for runtimes below 1 hour. We find that only 3 methods achieve satisfactory performance: the two methods introduced here, and the alignment-free method. The other two methods, Scoper and Partis, are limited to VJ*l* classes of small size (< 10^3^).

**FIG. 4.**
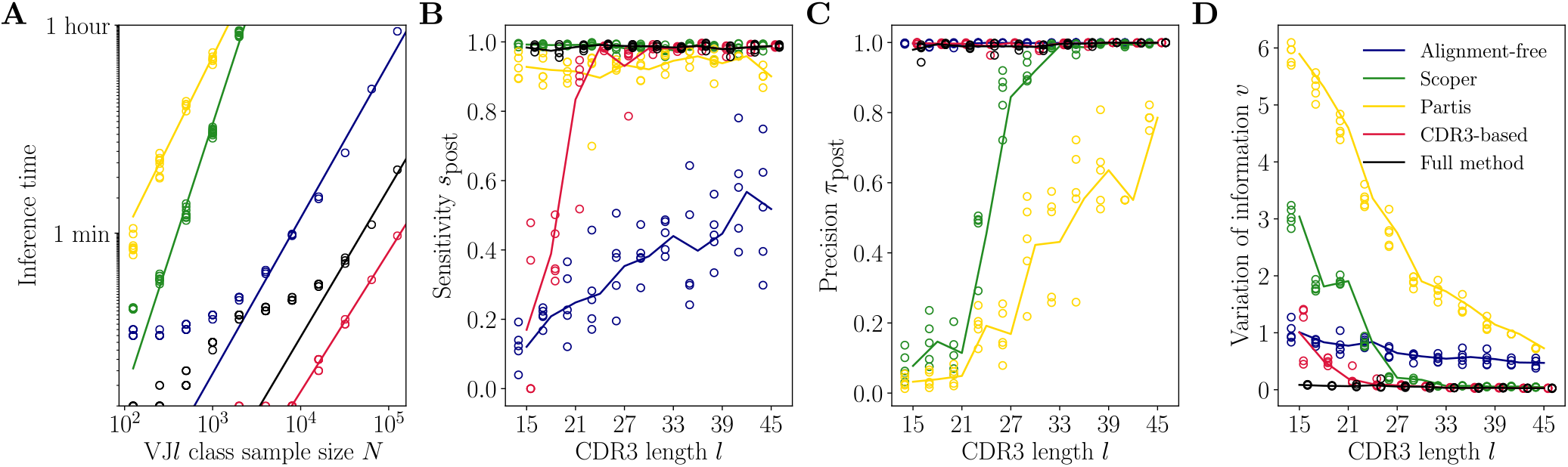
Benchmark of the alternative methods. (A) Comparison of inference time using subsamples from the largest VJ*l* class found in the donor 326651 of [1]. (B) Comparison of a posteriori sensitivity *s*_post_ across CDR3 length *l* in synthetic datasets. Solid lines represent the mean value averaged over 5 synthetic datasets. (C) Comparison of a posteriori precision *π*_post_ across CDR3 length *l* in synthetic datasets. (D) Variation of information *v* across CDR3 length *l* in synthetic datasets.

To compare the five algorithms in finite time, we test the accuracy of the methods using synthetic datasets with different CDR3 lengths. We focus on short CDR3s, *l* ∈ [15, 45], which are the most challenging for lineage inference. Each dataset contains 10^4^ unique sequences, so that the dominant VJ*l* class is typically of size ~ 10^3^ and can be handled by all 5 algorithms. We measure performance using three metrics applied to the resulting partition: pairwise sensitivity *s*_post_ (Fig. 4B), pairwise precision *π*_post_ (Fig. 4C), and the variation of information *v* (Fig. 4D). Pairwise sensitivity *s*_post_ and precision *π*_post_ are *a posteriori* analogs of the a priori estimates defined before in Eqs. 1 and 3, now computed after propagating links through the transitivity rule of single linkage clustering. Their value reflects not only the accuracy of the adaptive threshold but is also affected by the propagation of errors in single linkage clustering. Variation of information is a global metric of clustering performance which measures the loss of information from the true partition to the inferred one, and is equal to zero for perfect inference and positive otherwise (Methods, section IV G).

We find that, out of the five tested methods, only our method achieved both high sensitivity and high precision across all CDR3 lengths. Our CDR3-based method achieves high precision everywhere by construction, but only reached good sensitivity for CDR3 lengths 24 and above. The alignment-free method also achieves high precision everywhere, but with low sensitivity, meaning that it erroneously breaks up clonal families into smaller subsets. These three methods achieve good precision thanks to the use of a null model for the negative pairs. On the contrary, Scoper has excellent sensitivity everywhere but only achieves high precision for large lengths (*l* > 30), suggesting that it erroneously merges short-CDR3 clonal families. Likewise, Partis has good sensitivity but poor precision at all lengths, meaning that many clonal families are erroneously merged again.

The variation of information offers a useful summary of the performance. All methods perform worse for shorter CDR3s, except our full method, which is reliable everywhere. The CDR3-based method performs equally well, but only for CDR3 lengths 24 and above, while Scoper performs well only for lengths 30 and above.

We conclude that the full method should be chosen for its consistently high sensitivity, specificity, and speed. In the case of the largest datasets, the faster CDR3-based method is a useful alternative for long enough CDR3s.

### F. Inference of clonal families in a healthy repertoire

We next use our method to infer the clonal families of the heavy-chain IgG repertoires of healthy donors from [1]. Fig. 5 summarizes key properties of the inferred clonal families of donor 326651. We take advantage of the consistency of our method across CDR3 lengths, as evidenced by the benchmark, to study how the lineage structure changes with the CDR3 length. To this end, we divide the dataset into 9 quantiles, each containing between 8 and 12% of the total number of sequences (Fig. 5A, inset).

**FIG. 5.**
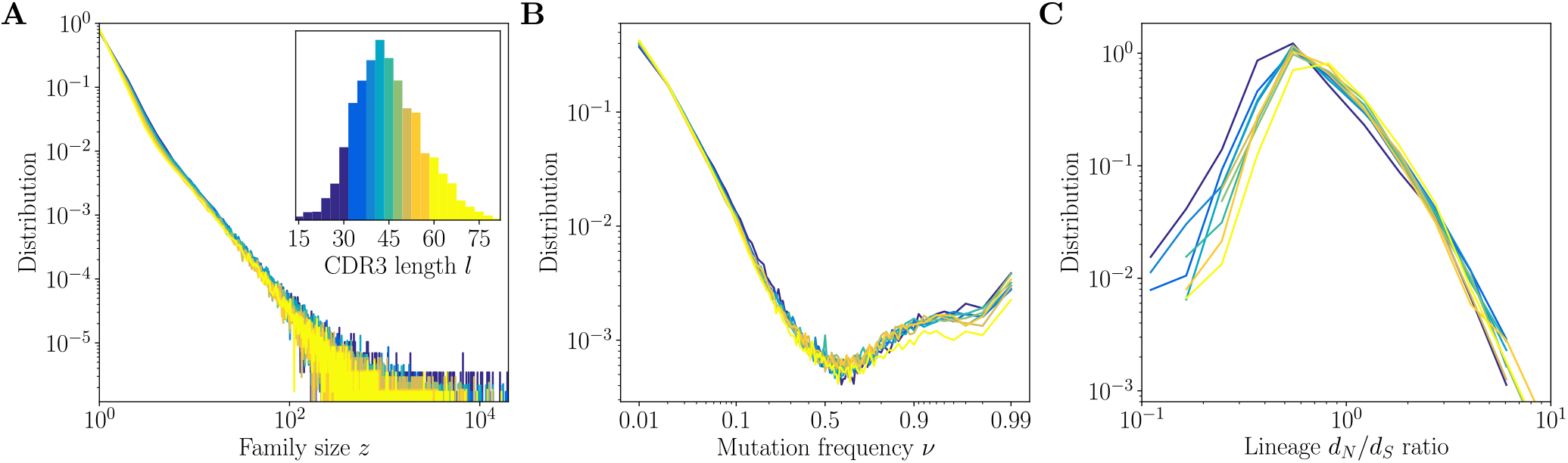
Inference results across CDR3 lengths. Inference results for donor 326651 of [1] are presented for 9-quantiles of the CDR3 distribution, each containing between 8 and 12% of the total number of sequences (corresponding to 9 colors in the inset of A). (A) Distributions of family size *z*. All CDR3 length quantiles exhibit universal power-law scaling with exponent −2.3 (B) Site frequency spectra estimated for families of sizes *z* = 100. Families of larger sizes were subsampled to *z* = 100 to subtract the influence of varying family sizes. (C) Distribution of lineage *d_N_/d_S_* ratios computed for polymorphisms in CDR3 regions over all lineages within each 9-quantile.

We find that across the 9 subsets of the data, the statistics of the lineage structure inferred with the mutations-based method are largely universal. The distribution of the clonal family sizes *z* (Fig. 5A) follows a power law across all CDR3 lengths under study, with no significant differences between different lengths. This results generalizes an earlier observation used above for generating synthetic datasets, but which was restricted to high-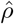, high-*l* VJ*l* classes, and justifies *a posteriori* the use of a universal power law in the generative model.

For the largest families, of size *z* ≥ 100, we compute two intra-lineage summary statistics: the site frequency spectrum, which gives the distribution of frequencies of point mutations within lineages, and the distribution of *d_N_/d_S_* ratios between non-synonymous and synonymous CDR3 polymorphisms within clonal families. To avoid the bias of the varying family sizes, we subsampled all families to size *z* = 100.

Under models of neutral evolution with fixed population size, the distribution of point-mutation frequencies *v* goes as *v*^−1^. Here we observe a non-neutral profile of the spectrum, with an upturn at large allele frequencies *v* > 0.5 (Fig. 5B). It is a known signature of selection or of rapid clonal expansion [23, 24]. We find that site frequency spectra are universal for all CDR3 lengths, suggesting that the dynamics that give rise to the structure of lineages and the subsequent dynamics that influence the sampling of family members, do not depend on the CDR3 length.

The lineage *d_N_/d_S_* ratio is also largely consistent across CDR3 lengths (Fig. 5C), while spanning 2 orders of magnitude, suggesting a wide gamut of selection forces. We could have expected longer loops to be under stronger purifying selection (lower *d_N_/d_S_*) to maintain their specificity and folding. Instead, we observe that short CDR3s have more lineages with low *d_N_/d_S_*. This may be due to different sequence context and codon composition in short versus long CDR3s. Short junctions are largely templated, whereas long junctions have long, non-templated insertions, and it was shown that templated regions have evolved their codons to minimize the possibility of non-synonymous mutations [25], which would lead to a lower *d_N_/d_S_*, regardless of selection.

## III. DISCUSSION

Clonal families are the building blocks of memory repertoire shaped by VDJ recombination and subsequent somatic hypermutations and selection. Repertoire sequencing datasets enable new approaches to understand these processes. They allow us to model the different sources of diversity and measure the selection pressures involved. To take full advantage of this opportunity we need to reliably identify independent lineages.

Here, we introduced a general framework for studying the methods for partitioning high-throughput sequencing of BCR repertoire datasets into clonal families. We have identified the main factors that influence the difficulty of this inference task: low clonality levels and short recombination junctions. We quantified the clonality level using the definition of pairwise prevalence *ρ* and introduced a method to estimate it a priori, without knowing the partition. We found the prevalence levels across VJ*l* classes to span three orders of magnitude (Figure 2B), unraveling the varying degree of complexity.

We leveraged the soNNia model of VDJ recombination to quantify the CDR3 diversity and constructed a null expectation for the divergence of independent recombination products. This null model enabled the design of a CDR3-based clustering method with an adaptive threshold that allows us to keep the precision of inference high across prevalences and CDR3 lengths. Owing to the prefix tree representation of the CDR3 sequences, this method is characterized by very short inference times thanks to avoiding all pairwise comparisons in single linkage clustering. As expected, we found that the adaptive threshold choice limits the sensitivity of inference in the regime of short junctions and low prevalence (Figure 3F).

To remedy the limitations of the CDR3-based approach we developed a mutations-based method. We found that including the phylogenetic signal of shared mutations in highly mutated sequences allows us to properly classify them into lineages despite significant CDR3 divergence. We studied the performance of the method using synthetic data and found significant improvement with respect to the CDR3-based approach: we extended the range of high-precision and high-sensitivity performance to cover all values of prevalence and CDR3 lengths observed in productive data (Figure 3G).

We have compared the two methods developed here with state-of-the-art approaches: the Partis [21] and Scoper [22] algorithms, and the alignment-free method [20]. We found that the crucial element to delivering a reliable partition is the ability to control for the false-positive rate across CDR3 lengths, as is the case of the alignment-free method, as well as the CDR-based and mutation-based methods introduced here. The need to estimate the false-positive rate is at the same time the main possible limitation of all of these approaches. In this work we took advantage of a soNNia model based on a neural network, and benefited from its expressivity to quantify the purifying selection that modifies the VDJ recombination statistics. We found the performance of this model satisfactory when applied to healthy memory repertoires, in agreement with previous findings [13, 19]. However, purifying selection is expected to be more pronounced in datasets of disease-specific cohorts and a default soNNia model may overestimate the diversity [26] and lead to underestimation of the fallout rate. The inference framework introduced here could still be applied with more sophisticated models of selection, and take advantage of higher levels of clonality that characterize many disease-specific datasets [10, 14].

We applied the mutations-based method to infer lineages in a repertoire of a healthy donor, sequenced at great depth [1]. We took advantage of the consistency our method exhibits across CDR3 lengths to find that the statistics of lineages, including a heavy-tail distribution of family sizes as well as signatures of selection, are universal and independent of the CDR3 length. This result implies that the dynamics of expansion, mutation, and selection are independent of the CDR3 and suggests they are dictated by the rules of affinity maturation and memory formation rather than BCR specificity. It advocates for the use of RNA sequencing data to quantify these general principles [26, 27]. Identifying clonal families with high accuracy is paramount in such approaches as it avoids the potential biases of different family sizes and varying levels of clonality.

The algorithm for clonal family identification presented here is a robust inference method that enables a reliable partition of a memory B-cell repertoire into independent lineages. Using synthetic datasets we demonstrated it is distinguished by consistently high precision and high sensitivity across different junction lengths and levels of clonality. It is therefore a useful tool to explore the diversity of the repertoires and improves our ability to interpret repertoire sequencing datasets.

## IV. METHODS

### A. Data preprocessing and alignment

We focus the analysis high-throughput RNA sequencing data of IgH-coding genes [1]. The sequences were barcoded with unique molecular identifiers (UMI) to correct for the PCR amplification bias. We aligned raw sequences using presto of the Immcantation pipeline [28] with tools allowing for correcting errors in UMIs and deal with insufficient UMI diversity. Reads were filtered for quality and paired using default presto parameters. We selected only sequences aligned with the IgG primer and therefore the lineage analysis is limited to the IgG subset of the repertoire. Pre-processed data was then aligned to V, D and J templates from IMGT [29] database using IgBlast [30].

Pairs of sequences stemming from the same VDJ recombination are expected to have the same CDR3 length *l* and align to the same V and J remplates. An exception could be caused by a insertion or deletion within the CDR3 that would alter its length as a result of the somatic hypermutation process. Such events are rare and generally selected against [31], therefore in what follows we shall assume the effect of these events is negligible. The inference could be also affected by the misalignment of either V or J templates but we previously found the effect of alignment errors to be insignificant for identifying VJ classes [32] (the alignment of the D template is error-prone and unreliable, hence not used in the inference procedure). Importantly, the two simplifications described here would result in decreased sensitivity of inference but are not expected to affect its precision.

### B. Modeling junctional diversity

The extraordinary diversity of VDJ rearrangments can be efficiently described and quantified using probabilistic models of the recombination process as well as subsequent purifying selection. Sequence-based models can assign to each receptor sequence s, its total probability of generation, *P*_gen_(*s*) [16, 33, 34] as well as a selection factor *Q*(*s*), inferred so as to match frequencies *P*_data_(*s*) of the sequences with a model-based distribution [19, 35, 36]

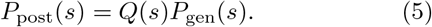

The *P*_gen_ model was inferred using unmutated out-of-frame sequences from [1] using the IGoR software [34]. The selection function *Q* model was learned using unmutated productive IgM sequences from [1] using the soN-Nia software [19].

The post-selection distribution *P*_post_ describes the diversity of the CDR3 regions and in doing so provides an expectation of pairwise distances between unrelated, independently generated sequences of same length *l* [19]. We can define

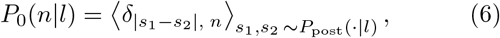

where |*s*_1_ – *s*_2_| stands for (Hamming) distance between sequences *s*_1_ and *s*_2_. This definition of the null distribution is a straightforward recipe for its estimation using (Monte Carlo) samples from *P*_post_.

Should *P*_post_ differ significantly from the empirical frequencies *P*_data_ one can resolve to the following alternative

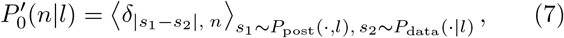

the equivalent of the negation distribution as defined in [20] and used in our evaluation of the alignment-free method [20] in the method benchmark analysis.

### C. Estimation of pairwise prevalence

Pairwise prevalence is defined as the ratio of pairs of related sequences to the total number of pairs of sequences in a given set. Related sequences share an ancestor and have diverged by independent somatic mutations, post-recombination. Low prevalence can be a major difficulty for any inference procedure as any misassignment (or fallout) will result in a drastic loss of sensitivity or precision. It is instrumental to have an a priori estimate of pairwise prevalence before the families are identified.

To estimate the prevalence from the distribution of distances *P*(*n*) for a given set of sequences (typically a VJ*l* class or *l* class) we propose the following expectation-maximization procedure. We stipulate the distribution in question is a mixture distribution of two components, *P*_0_(*n*), the expectation for unrelated sequences defined as above, and *P*_1_(*n*), describing related sequences, modeled using a Poisson distribution

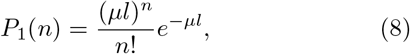

where *μ* is the mean divergence per basepair. In case of *l* class comprising very many VJ*l* classes the resultant shape of the positive distribution is often closer to a geometric profile, and is then modeled using *P*_1_(*n*) = (1 – *M*)*M^n^*, where 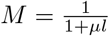. In sum

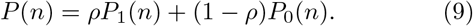

In a standard fashion, we proceed iteratively by calculating the expected value of the log-likelihood (pairs of sequences indexed by *i*)

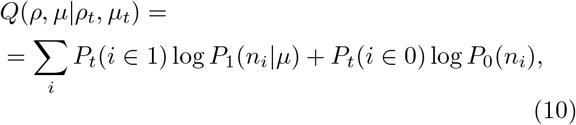

where the membership probabilities are defined as

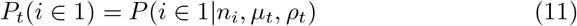

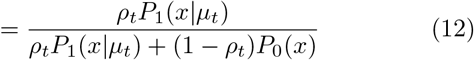

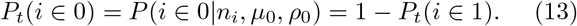

We then find the maximum

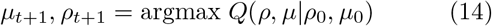

and iterate the expectation and maximization steps until convergence, |*ρ*_*t*+1_ – *ρ_t_*| < *ϵ*, to obtain 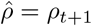.

Results for largest VJ*l* class within each *l* class can be found in Fig. S3 and results for *l* classes using a geometric distribution can be found in Fig. S6. Dependence of maximum likelihood prevalence estimates 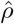 on class size *N* is plotted in Fig. S7.

### D. CDR3-based clustering

The standard method for CDR3-based inference of lineages proceeds through single-linkage clustering with a fixed threshold on Hamming distance divergence [12, 24, 37, 38]. This crude method suffers from inaccuracy as it loses precision in the case of highly-mutated sequences and junctions of short length. However, if junctions are stored in a prefix tree data structure [39] single-linkage clustering can be performed without comparing all pairs and hence is typically orders of magnitude faster than alternatives. The prefix tree is a search tree constructed such that all children of a given node have a common prefix, the root of the tree corresponding to an empty string, and leaves corresponding to unique sequences to be clustered. To find neighbors of a given sequence it suffices to traverse the prefix tree from the corresponding leaf upwards and compute the Hamming distance at branchings. This method limits the number of unnecessary comparisons and greatly improves the speed of Hamming distance-based clustering [40]. We implement the prefix tree structure to accommodate CDR3 sequences. Briefly, all the CDR3 sequences of identical length are stored in the leaves of a prefix tree [40, 41], implemented as a quaternary tree where each edge is labeled by a nucleobase (A, T, C, or G). The neighbors of a specific sequence are found by traversing the tree from top to bottom, exploring only the branches that are under a given Hamming distance from the sequence. Clusters are obtained by iterating this procedure and removing all the neighbors from the prefix tree until no sequences remain. The package is coded in C++ with a Python interface and is available independently. The time performance of this method for high-sensitivity and high-specificity partitions is studied as a part of the method benchmark analysis.

We take advantage of the speed of a prefix tree-based clustering and perform single-linkage clustering with an adaptive threshold. For any dataset, we define two CDR3-based partitions: high-sensitivity and high-precision clustering.

To ensure high sensitivity *s*\*, the threshold *n* of clustering should grow linearly with the CDR3 length *l*,

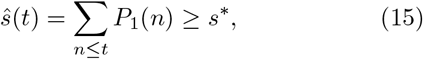

where we assume the Poisson or geometric distribution of positives. We find effectively 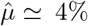, so that the threshold for high-sensitivity clustering reads *t*_sens_ = ⎣10%*l*⎦ +1 for *s* ≃ 95% and *t*_sens_ = ⎣15%*l*⎦ + 1 for *s* ≃ 99% (where ⎣.⎦ denotes the floor function).

This is to be juxtaposed with a high-precision partition, where the fall-out rate 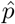 (1 – specificity) is adjusted using the estimate of pairwise prevalence 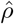 and the *P*_post_ prediction for the null distances distribution *P*_0_(*n|l*). On one hand the fallout rate can be computed as

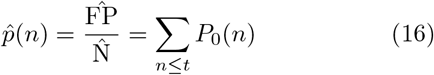

and on the other, by inverting the relation (1) for required precision *π*\*,

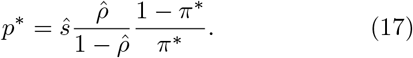

Given a required precision *π*\* we can now find the corresponding fallout rate *p*\* and the high-precision Hamming distance threshold *t*\* so that 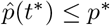.

Finally, the structure of families leads to propagation of errors that lowers the precision with respect to the a priori estimate 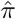. Denoting family size as *z*, one error accounted for in 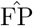 causes, on average, 〈*z*〉^2^ – 1 extra errors by merging two families. If the a priori precision 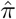 is high, we can neglect the second order effect of these two families simultaneously affected by other 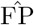 pairs). Therefore the expected precision (1) of the resulting partition reads

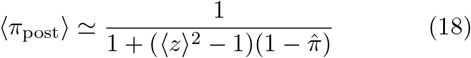

where we assumed *ŝ* ≃ 1. For 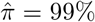 and 〈*z*〉 ≃ 2 this formula gives 〈*π*_post_) ≃ 97%.

### E. Synthetic data generation

To generate synthetic data we make use of the lineages identified in the high-sensitivity and high-precision regime of CDR3-based inference (Fig. 3F), we denote the set of these lineages by 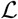. We assume that to good approximation the mutational process and the selection forces that shaped the mutational landscape in these lineages do not depend on the CDR3 length.

To test the performance of different inference methods across CDR3 lengths, we build synthetic datasets of fixed length.

In the first step, we choose the number of families *N*. We then draw *N* independent family sizes from the family size distribution of the form observed in healthy datasets

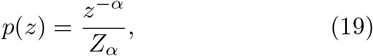

where *Z_α_* = ∑_*z*≥1_*z*^−*α*^ = *ζ*(*α*, 1). In the next step, we assign a naive progenitor to each lineage by sampling from the *P*_post_ distribution, selecting sequences with a prescribed length *l* (Fig. S4). We then choose a lineage in the set of reconstructed lineages 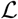 at random amongst lineages of size *z* (or, for large sizes, the lineage of the closest size smaller than *z*). We then identify all unique mutations in the true lineage and for each mutation denote the labels of members of the lineage that carry it. For each mutation, this defines a configuration of labels, one of 2^*z*^ – 1 possible. We subsequently loop through observed configurations and choose new positions for all mutations to apply them to the synthetic progenitors of the ancestor, using the position- and context-dependent model of Ref. [32]. The number of mutations assigned to a given configuration is rescaled by a factor 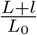 where *L* is the templated length of the synthetic ancestral sequence and *L*_0_ is the templated length of the model lineage from 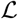.

This way a synthetic lineage preserves all properties of the lineages of long CDR3s found in the data, particularly the mutational spectra (Fig. S5) except for the ancestral sequences and the identity of mutations.

### F. Mutations-based method

We compute the expected distributions of the CDR3 Hamming distance *n*, and the number of shared mutations *n*_0_, under a uniform mutation rate assumption. In other words, we assume that the probability that a given position was mutated, given a mutation happened somewhere in a sequence of length *L*, equals *L*^−1^. It follows that the probability that a given position has not mutated once in a series of *n* mutations is (1 – *L*^−1^)^*n*^.

#### Expectation of *n*_0_ under the null hypothesis

For *n*_0_ shared mutations, under the null hypothesis (we operate under the null hypothesis here since otherwise to estimate *n*_0_ we would need to make assumptions about the law that governs B-cell phylogeny topologies), the likelihood reads

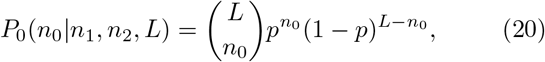

where the probability that the same position independently mutated in series of *n*_1_ and *n*_2_ mutations is

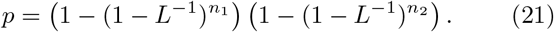

In the limit of large *L*,

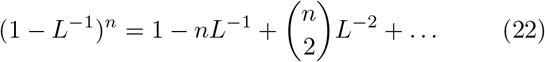

and

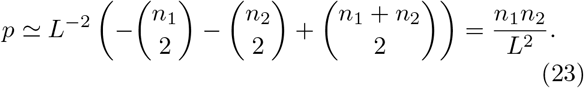

Hence

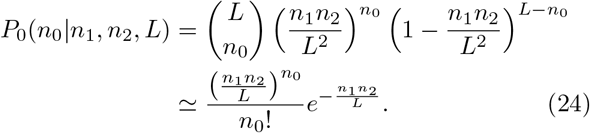

The characteristic scale is then the fraction 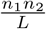, and

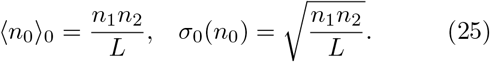

#### Expectation of *n* under the hypothesis of related sequences

The *n* divergence of two CDR3s is interpreted as divergent mutations under the hypothesis that s1 and s2 are related. These mutations were harbored in parallel with *n_L_* = *n*_1_ + *n*_2_ – 2*n*_0_ mutations that occurred in the templated regions (*n*_0_ mutations arrived before the divergence of the two sequences began).

Under the assumption of a uniform mutation rate, the *n_L_* mutations inform the prediction of the number of mutations expected in the CDR3. Indeed, they are related through a hidden variable, the expected number of mutations per base pair, denoted *μ*. Integrating over this quantity we obtain

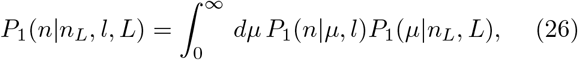

where we convolute the positive distribution (8),

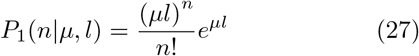

and, using the Bayes rule under uniform prior over *μ*,

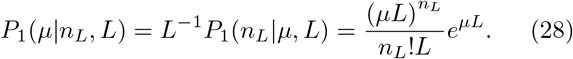

The result is a negative binomial distribution,

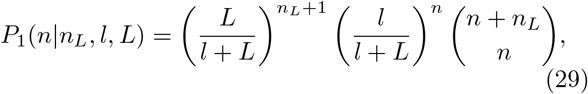

with

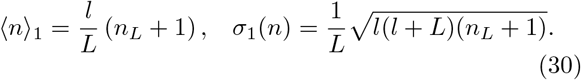

The mutations-based method relies on the results (30) and (25) to define the rescaled variables (4)

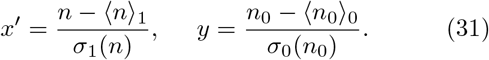

We perform single linkage clustering with the distance between two sequences defined as *x′* – *y* and imposing a threshold *x′* – *y* ≤ *t* where *t′* is chosen based on the prevalence, analogically to adaptive threshold choice in the CDR3-based inference method. To this end we use soNNia-based estimate of null distribution *P*_0_(*n|l*) (6), the data-derived distribution of the number of mutations, *P*(*n*_1_), and further assume 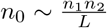 to compute the new null distribution *P*_0_(*x′* – *y|l*).

### G. Evaluation methods

In this section, we introduce the variation of information *υ*, used for evaluating alternative methods for clonal family inference in the benchmark analysis. It is a useful summary statistic to quantify the performance of inference as it is affected by its precision as well as sensitivity [42]. Variation of information *υ*(*r*,*r*\*) measures the information loss from the true partition *r*\* to the inference result *r* [43, 44]. To define the variation of information we first introduce the entropy *S*(*r*) of a partition *r* of *N* sequences into clusters *c* as

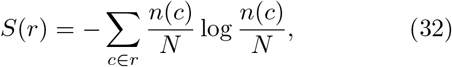

where *n*(*c*) denotes the number of sequences in cluster *c*. The mutual information between two partitions *r* and *r*\* can then be computed as

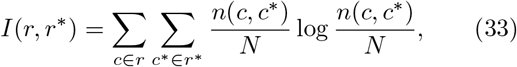

where *n*(*c*, *c*\*) denotes the number of overlapping elements between cluster *c* in partition *r* and cluster *c*\* in partition *r*\*. Finally, variation of information is given by

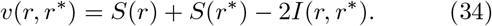

Variation of information is a metric in the space of possible paritions since it is non-negative, *υ*(*r*, *r*\*) ≥ 0, symmetric, *υ*(*r*, *r*\*) = *υ*(*r*\*, *r*), and obeys the triangle inequality, *υ*(*r*_1_, *r*_3_) ≤ *υ*(*r*_1_, *r*_2_) + *υ*(*r*_2_, *r*_3_) for any 3 partitions [43].

### H. Code and data availability

The HILARy tool with Python implementations of the CDR3 and mutations-based methods introduced above can be found at github.com/statbiophys/HILARy. The standalone prefix tree implementation can be found at github.com/statbiophys/ATrieGC.

## ACKNOWLEDGEMENTS

The study was supported by European Research Council COG 724208 and ANR-19-CE45-0018 ‘RESP-REP’ from the Agence Nationale de la Recherche grants and DFG grant CRC 1310 ‘Predictability in Evolution’.

## V. SUPPLEMENTARY INFORMATION

**FIG. S1.**
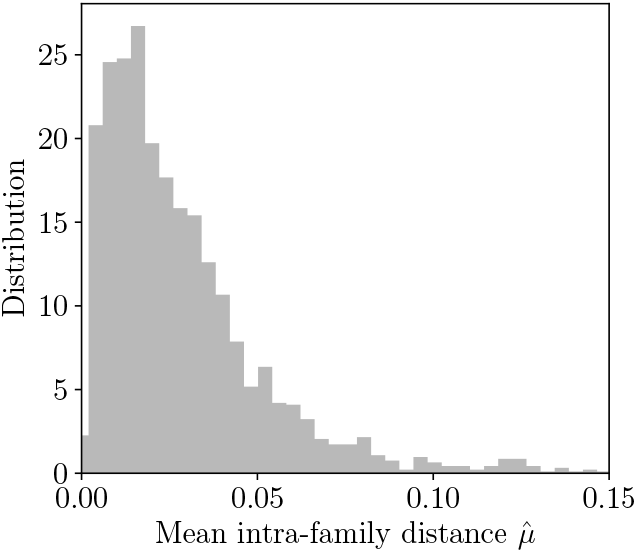
Distribution of the maximum likelihood estimates of mean intra-family distance 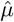 across VJ*l* classes.

**FIG. S2.**
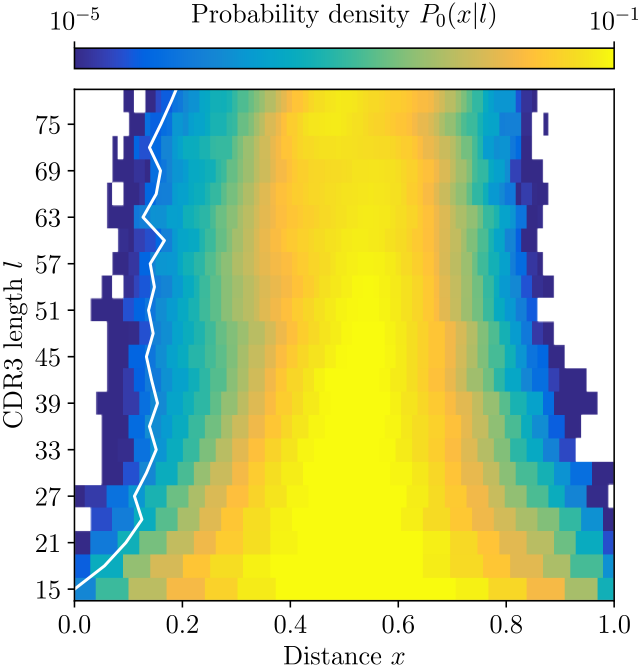
Null distribution *P*_0_(*x|l*) of CDR3 distances between unrelated sequences for *l* ∈ [15, 81], computed by soNNia software. White line denotes a growing threshold ensuring a fallout rate *p* < 10^−4^ as determined by this distribution.

**FIG. S3.**
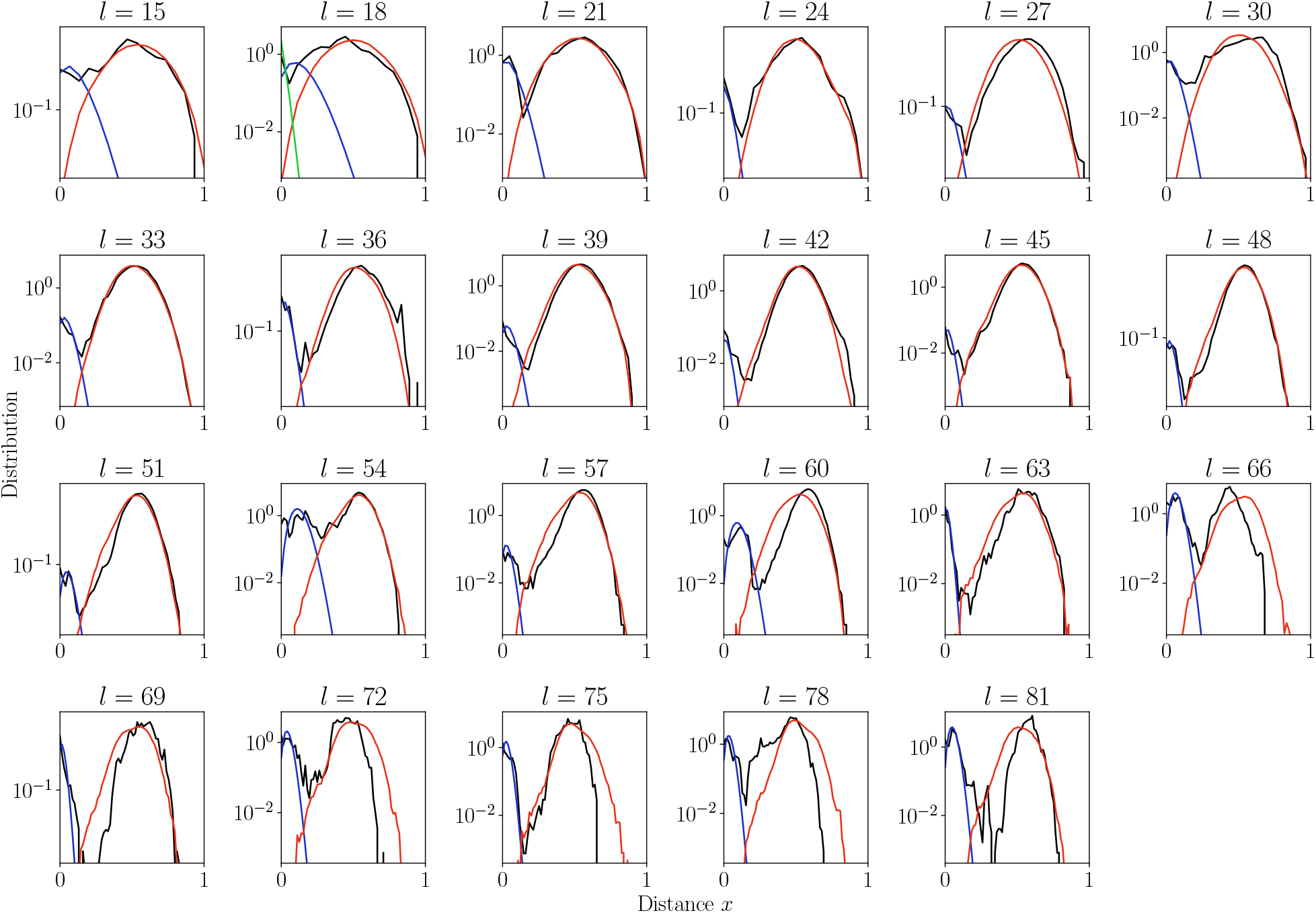
Distribution of normalized Hamming distances, *x* = *n/l*, for largest VJ*l* class for each CDR3 length *l* = 15,†, 81 (black). We fit the distribution by a mixture of positive pairs (*P*_1_(*x*|*μ*) in blue), and negative pairs (*P*_0_(*x*), in red). For *l* = 18 the estimate 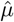 is too large results in large fitting error and for sensitivity computation we used global *μ* = 4% (in green).

**FIG. S4.**
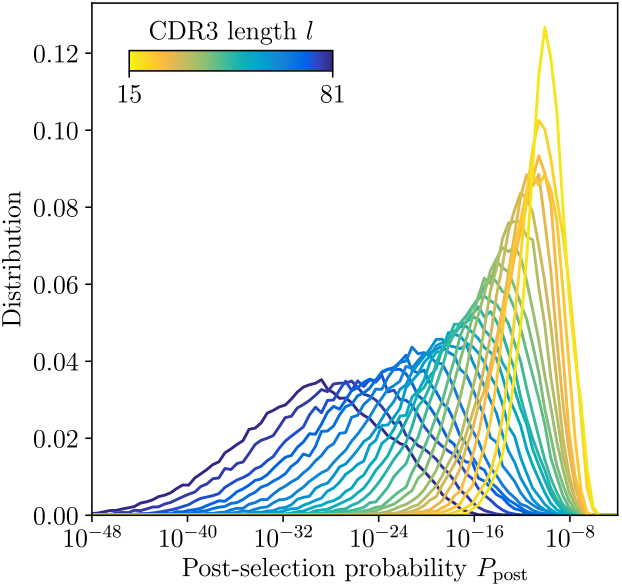
Distribution of post-selection probabilities *P*_post_ of CDR3 nucleotide sequences computed using soNNia across CDR3 lengths. Short junctions are on average more likely to be generated in VDJ recombination and pass subsequent selection [19]. This makes inference in low-*l* classes more difficult, a feature reflected by synthetic dataset constructed by sampling unmutated lineage progenitors from the soNNia model.

**FIG. S5.**
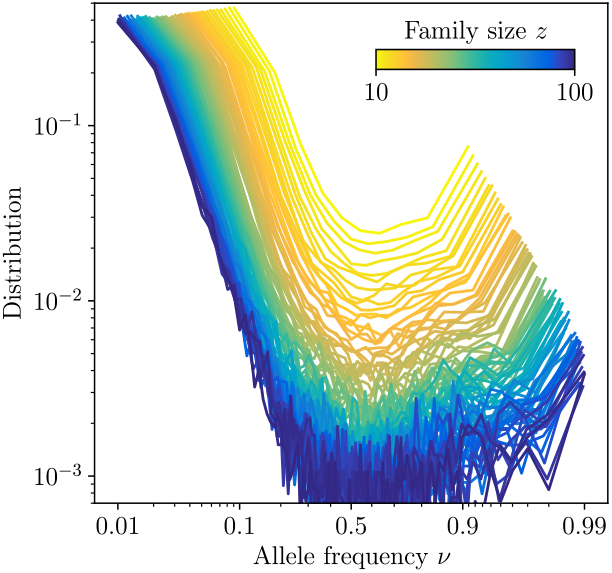
Site frequency spectra estimated for families identified using high-precision CDR3-based inference method in the subset of the data where this approach is highly reliable (large-*l* and large-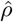 regime). The distributions are shown for families of varying family size, *z* ∈ [10, 100] and averaged over all families of a given size. Together with the exact configuration of sequences carrying a given substitution, synthetic datasets of the same signatures of mutations and clonal expansions can be generated.

**FIG. S6.**
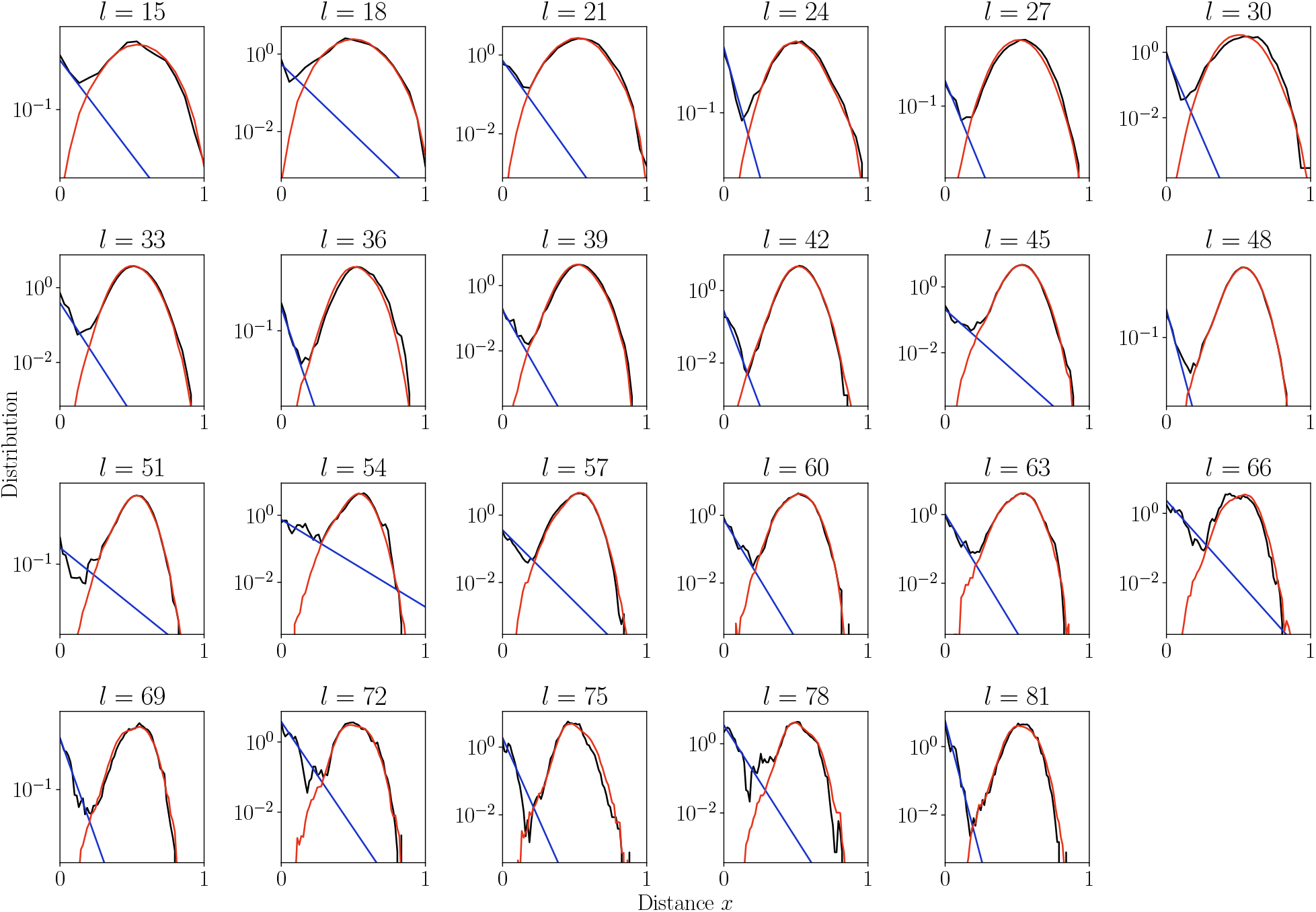
Distribution of normalized Hamming distances, *x* = *n/l*, for *l* classes, averaging over all VJ*l* classes. We fit the distribution by a mixture of positive pairs using a geometric distribution (*P*_1_(*x*|*μ*) in blue), and negative pairs (*P*_0_(*x*), in red). The corresponding prevalence estimates 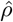 are used for small VJ*l* classes for which this parameter cannot be reliably estimated independently.

**FIG. S7.**
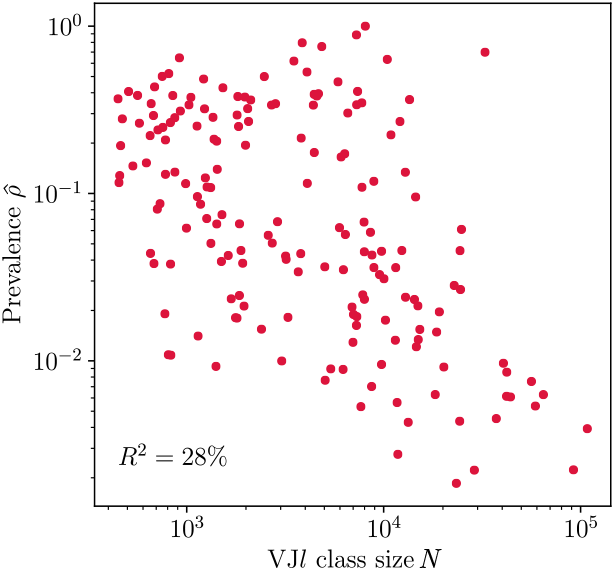
Dependence of prevalence estimates 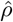 on VJ*l* class size *N* for largest classes in donor 326651 from [1]. 28% of variation in prevalence estimates can be explained by variation in VJ*l* class sizes.

## Notes

### Competing Interest Statement

The authors have declared no competing interest.

